# Identification of a blood test-based biomarker of aging through deep learning of aging trajectories in large phenotypic datasets of mice

**DOI:** 10.1101/2020.01.23.917286

**Authors:** Konstantin Avchaciov, Marina P. Antoch, Ekaterina L. Andrianova, Andrei E. Tarkhov, Leonid I. Menshikov, Olga Burmistrova, Andrei V. Gudkov, Peter O. Fedichev

## Abstract

We proposed and characterized a novel biomarker of aging and frailty in mice trained from the large set of the most conventional, easily measured blood parameters such as Complete Blood Counts (CBC) from the open-access Mouse Phenome Database (MPD). Instead of postulating the existence of an aging clock associated with any particular subsystem of an aging organism, we assumed that aging arises cooperatively from positive feedback loops spanning across physiological compartments and leading to an organism-level instability of the underlying regulatory network. To analyze the data, we employed a deep artificial neural network including auto-encoder (AE) and auto-regression (AR) components. The AE was used for dimensionality reduction and denoising the data. The AR was used to describe the dynamics of an individual mouse’s health state by means of stochastic evolution of a single organism state variable, the “dynamic frailty index” (dFI), that is the linear combination of the latent AE features and has the meaning of the total number of regulatory abnormalities developed up to the point of the measurement or, more formally, the order parameter associated with the instability. We used neither the chronological age nor the remaining lifespan of the animals while training the model. Nevertheless, dFI fully described aging on the organism level, that is it increased exponentially with age and predicted remaining lifespan. Notably, dFI correlated strongly with multiple hallmarks of aging such as physiological frailty index, indications of physical decline, molecular markers of inflammation and accumulation of senescent cells. The dynamic nature of dFI was demonstrated in mice subjected to aging acceleration by placement on a high-fat diet and aging deceleration by treatment with rapamycin.

## I. INTRODUCTION

An ever-increasing number of physiological state variables, such as blood cell counts and blood chemistry [1–4], DNA methylation [4–8], locomotor activity [4, 9–11], and exploratory behavior [9, 12], have been investigated in association with aging and used to quantify aging progression in in-vivo experiments and in future anti-aging clinical trials [4]. Most of the common statistical models used in aging studies require chronological age or age at death as labels for training; however, this data are rarely available in sufficient quantity from human and laboratory animal cohorts. Even less information is available regarding the change in biomarkers of aging and frailty across the lifespan of individual animals or patients in response to lifespan-modifying interventions. Thus, the field would benefit from development of a convincingly justified easily measurable and reliable biomarker of aging ideally obtainable from conventional and automated measurements such as routine blood tests.

To produce a quantitative description of aging process in mice, we turned to the largest open-access source of phenotypic data, the Mouse Phenome Database (MPD) [13, 14]. The MPD contains a wide range of phenotype data sets including behavioral, morphological and physiological characteristics and involving a diverse set of inbred mouse strains. In the present work, we implemented biomarker of aging based on complete blood cell (CBC) measurements. CBC test is a well established, easily obtainable laboratory analysis protocol and it has a long list of applications in both clinical medicine and biomedical research [15].

Principal Components Analysis (PCA) revealed that the fluctuations in CBC variables in the MPD are dominated by the dynamics of a single cluster of features, jointly deviating from the initial state and increasing in variance with age.

The behaviour is typical for non-equilibrium complex systems with strong interactions between the components operating close to the critical or tipping point separating the stable and the unstable regimes [16]. Under the circumstances, the organism state fluctuations should be driven by the dynamics of a single variable, that is the organism-level property having the meaning of the order parameter corresponding to the unstable phase [17] and associated with aging drift and mortality acceleration [18].

To generate the biomarker of aging, we built a state-of-the-art deep artificial neural network composed of de-noising autoencoder (AE) and the auto-regressive (AR) model, that is a computational metaphor for the dynamics of the order parameter. The network output variable exhibited the most desirable properties of a biological age marker: it increased exponentially with age, predicted the remaining lifespan of the animals, correlated with multiple hallmarks of aging, and was henceforth referred to as the dynamic frailty index (dFI). The dynamic character of dFI was demonstrated in experiments involving treatments previously shown to accelerate (high-fat diet) or decelerate (rapamycin) aging in mice.

Therefore, we conclude that dFI is an accurate, easily accessible biological age proxy for experimental characterization of aging process and anti-aging interventions. On a more conceptual level, our work demonstrates that the auto-regressive analysis provided by the AE-AR deep learning architecture may be a useful tool for the fully unsupervised (label-free) discovery of biological age markers from any type of phenotypic data involving longitudinal measurements.

## II. RESULTS

### A. Overview of aging in the Mouse Phenome Database

We started by building a training set from the largest publicly available source of phenotypic data, the Mouse Phenome Database (MPD) [13, 14]. To achieve the best possible compatibility with earlier studies, we scanned the database records to maximize the number of available measurements common to those used in the construction of the physiological frailty index (PFI) in [19]. As a result, we chose a subset of twelve complete blood count (CBC) measurements from nine datasets, altogether including more than 7, 500 animals (see Table S2 for a complete list of datasets used for training of the models in this work).

To visualize the 12-dimensional CBC data from the MPD, we performed principal component analysis (PCA), that is a computational technique commonly used for multivariate data analysis [20–22]. PCA of the MPD slice representing fully-grown animals (exceeding the age of 25 weeks old) turned out to be particularly simple, see Fig. 1**a**. In this case, most of the variance in the data (27%) is explained by the first PC score, *z*_0_, which is strongly associated with age. None of the subsequent PC scores (*z*_1_, *z*_2_, etc., each explaining 20%, 16%, etc. of the data variance, respectively) showed any reasonable correlation with age. However, each of the PC scores was associated with a distinct cluster of biologically related blood features, as shown in Fig. 1**b**. For example, the first two PC scores could be predominantly connected with the red and white blood cell counts, respectively.

**FIG. 1:**
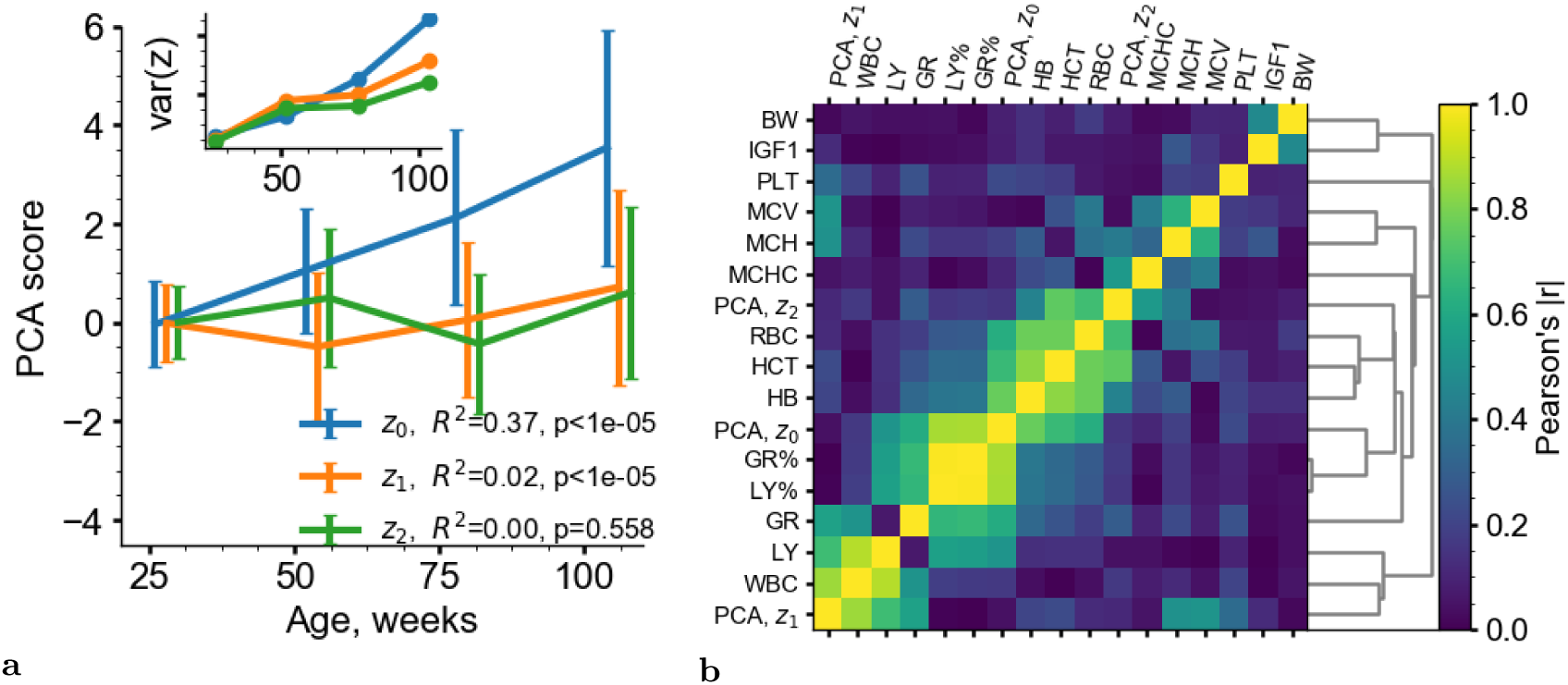
Principal Component Analysis (PCA) of the MPD slice corresponding to mature animals (age greater than 25 weeks). **a** The graphs represent the average of the PC scores in subsequent age groups (the error bars are at the standard deviation). The inset shows that the variance for all PC scores increase with age. **b** Clustering of CBC features and PC scores in the training dataset. The colors represent the Pearson’s correlation coefficient (absolute value) as indicated by the scale on the right side of the figure.

Notably, the largest variance in the data representing the full dataset was rather associated with animal growth and maturation. The first PC score is associated with the age of animals younger than 25 weeks old but does not change substantially after that age, see Fig. S1. The second, PC score exhibited an association with age over the entire available age range. This means that aging and early development in mice are different phenotypes and henceforth we perform all our calculations using the data from animals aged older than 25 weeks.

The first PC score, *z*_0_, was the only PCA variable associated with the remaining lifespan of the animals. This was determined by using Spearman’s rank-order correlation tests to evaluate potential associations between the first three PC scores and the age at death within cohorts of mice of the same age and sex (Table I). The *z*_0_ variable had therefore the most desirable properties of biological age.

**TABLE I:**
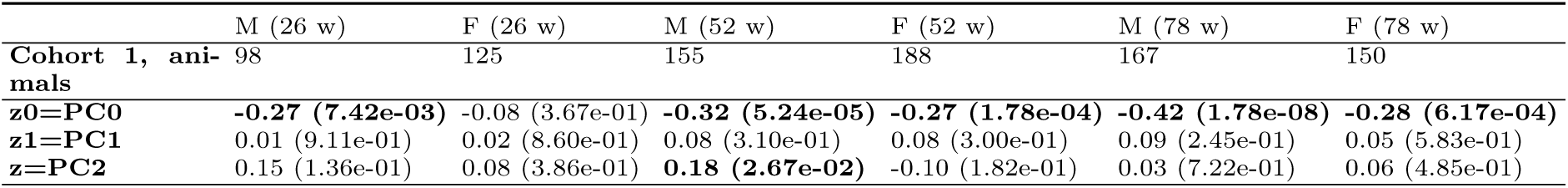
Spearmans rank-order correlation values and the corresponding p-values (in parentheses) for the top PC scores with the remaining) lifespan (the significant correlations (*p* < 0.05) are highlighted in bold.

Variance of the PCA scores and hence the biological age also grew with age (see the inset in Fig. 1**a**), which is a signature of stochastic broadening. The dimensionality reduction revealed by the PCA and the association of the large-scale fluctuations driving the slow evolution or disintegration of the system are characteristic of criticality, which is a special case of the dynamics of a complex system unfolding near a bifurcation or a tipping point, on the boundary of a dynamic stability region [16, 23].

### B. Aging, critical dynamics of the organism’s state and the dynamic frailty index (dFI)

The dynamics of the order parameter associated with the unstable phase is a measure of the aging drift and mortality acceleration in aging organisms [18] and henceforth is to be referred to as the dynamic frailty index (dFI). In this section, we summarize the necessary theoretical framework required for identification of the biomarker and quantitative description of aging in biological data.

Over sufficiently long time-scales, the fluctuations of physiological indices (such as CBC features), *x*_*i*_, are expected to follow the dynamics of the order parameter, *z* = dFI: *x*_*i*_ = *b*_*i*_*z* + *ξ*_*i*_. Here *ξ*_*i*_ is noise, *b*_*i*_ is a vector that may differ across species, and the integer index *i* enumerates the measured features.

Close to the tipping point, the dynamics of the physiological state is slow and hence the variable *z* satisfies the stochastic Langevin equation with the higher order time derivative terms neglected:

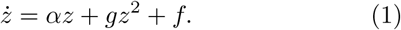

Here the linear term, *αz*, on the right side of the equation represents the effect of the regulatory network stiffness governing the responses of the organism to small stresses producing small deviations of the organism state from its most stable position. The following term, *gz*^2^, represents the lowest order non-linear coupling effects of regulatory interactions.

The stochastic forces *f* represent external stresses and the effects of endogenous factors not described by the effective Eq. 1. Naturally, we assume that random perturbations of the organism state are serially uncorrelated, so that ⟨*f* (*t*)*f* (*t*′)⟩ ∼ *B*, where *B* is the power of the noise, and ⟨…⟩ stands for averaging along the aging trajectory.

The equation establishes the “law of motion” for the organism’s physiological state. It is a mathematical relation between the rate of change of the organism state variable, *ż* = *dz/dt*, on the left side of the equation, and the effects of deterministic (*αz, gz*^2^) and stochastic forces (*f*), on the right side.

Depending on the sign of the stiffness coefficient, *α*, the organism state may be dynamically stable (if *α* < 0) or unstable (if *α* > 0). In the latter case, small deviations of the organism state get amplified over time so that no equilibrium is possible and the solution of Eq.1 describes an aging organism. Typically, *α* is small, and hence, the evolution of the physiological indices exhibits hallmarks of critically: it is slow (critical slowing down) and the fluctuations of the physiological state following the variations in *z* are large [*z*^2^ ∼ *B* exp(2*αt*)*/*2*α* (critical fluctuations).

Very early in life, the deviations from the critical point are small and the evolution of the organism state is dominated by diffusion. Later in life, the linear term takes over such that the deviations from the youthful state accelerate exponentially:

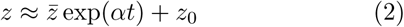

where 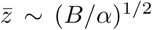 and *z*_0_ are constants representing the accumulated early effects of random and deterministic forces, respectively.

Finally, once dFI is sufficiently large, *z* ≳ *Z* = *α/g*, the non-linear terms take over, disintegration of the organism state proceeds at a rate greater than exponential, and the animal dies in a finite time. Mortality in this model increases up to the average lifespan 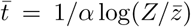. Mortality is a complex function of the order parameter *z* and hence of the chronological age. The mortality acceleration rate at the age corresponding to the average lifespan is of the same order of *α*.

### C. Identification of dFI from longitudinal data by applying a deep neural network

To identify the dFI from CBC measurements we performed a fit of the experimental data from MPD onto solutions of Eq. 1 with the help of an artificial neuron network. Altogether we used 7616 samples from 9 MPD datasets as the training set (see Material and Methods and Table S2). We employed a combination of a deep auto-encoder (AE) and a simple auto-regression (AR) model for modal analysis (AE-AR; see neural network architecture in Fig. 8). At its bottleneck, the encoder arm of the AE produced a compressed 4-dimensional representation **y** of the input, the 12-dimensional physiological state vectors **x** built from the available CBC measurements. The decoder arm reconstructed the original 12-dimensional state 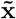 from the bottle-neck features.

**FIG. 2:**
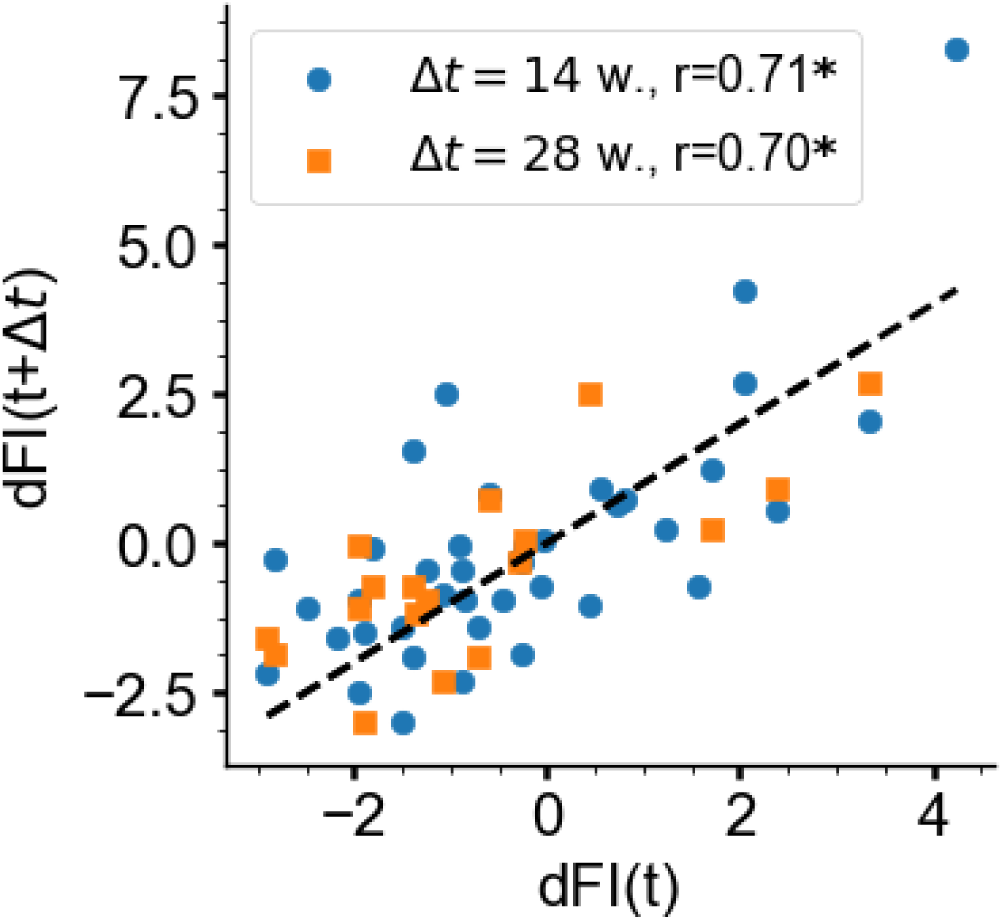
Auto-correlation properties of age-adjusted dFI across sampling intervals Δ*t* of 14 (blue circles) and 28 (orange squares) weeks. (* marks statistically significant correlations, *p* < 0.001)

**FIG. 3:**
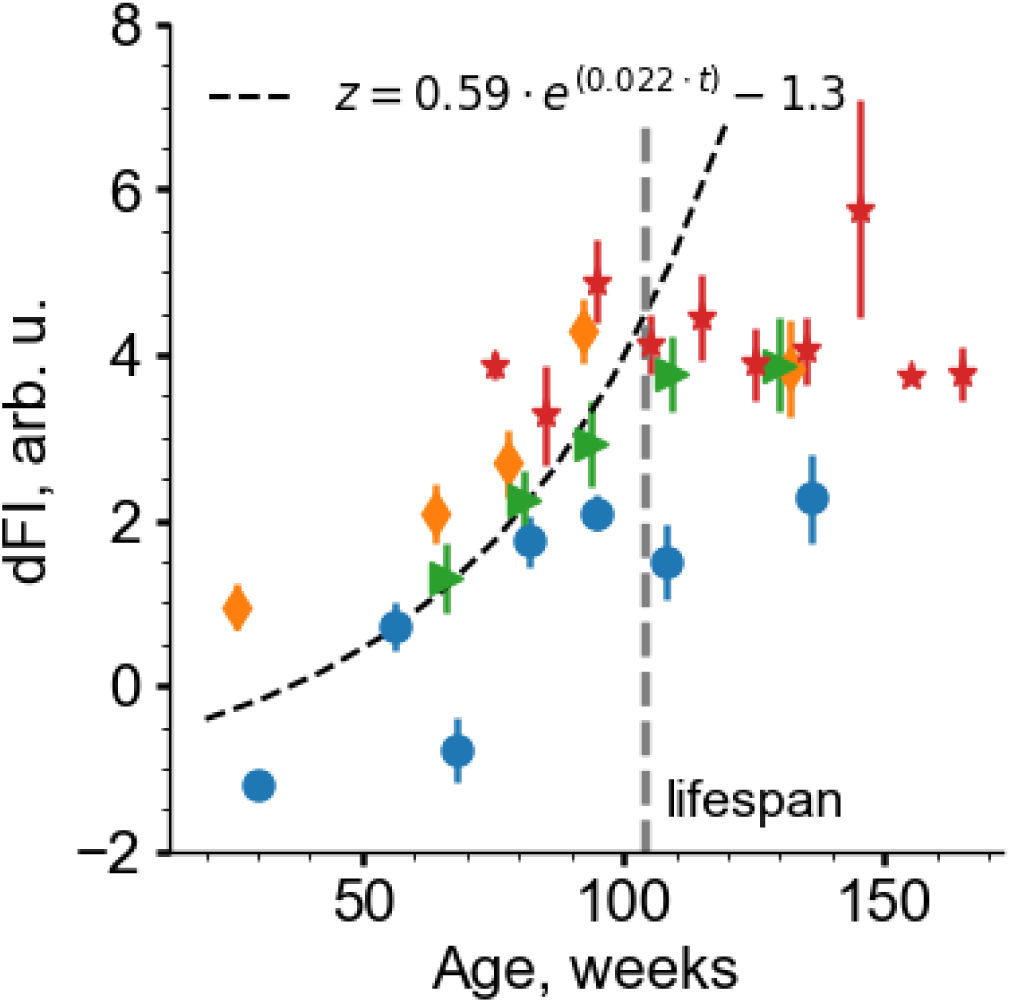
Dynamical frailty index (dFI) as a function of age in the test experiments: MA0071 (males, orange diamonds), MA0071 (females, blue circles) and MA0072 (green triangles). The black curved dashed line is the exponential fit in the age groups younger than the average lifespan of NIH Swiss mice (indicated by the dashed vertical line). Red stars mark the average dFI in age-matched groups of frail animals from the MA0073 cohort. All data are presented as mean±SEM.

**FIG. 4:**
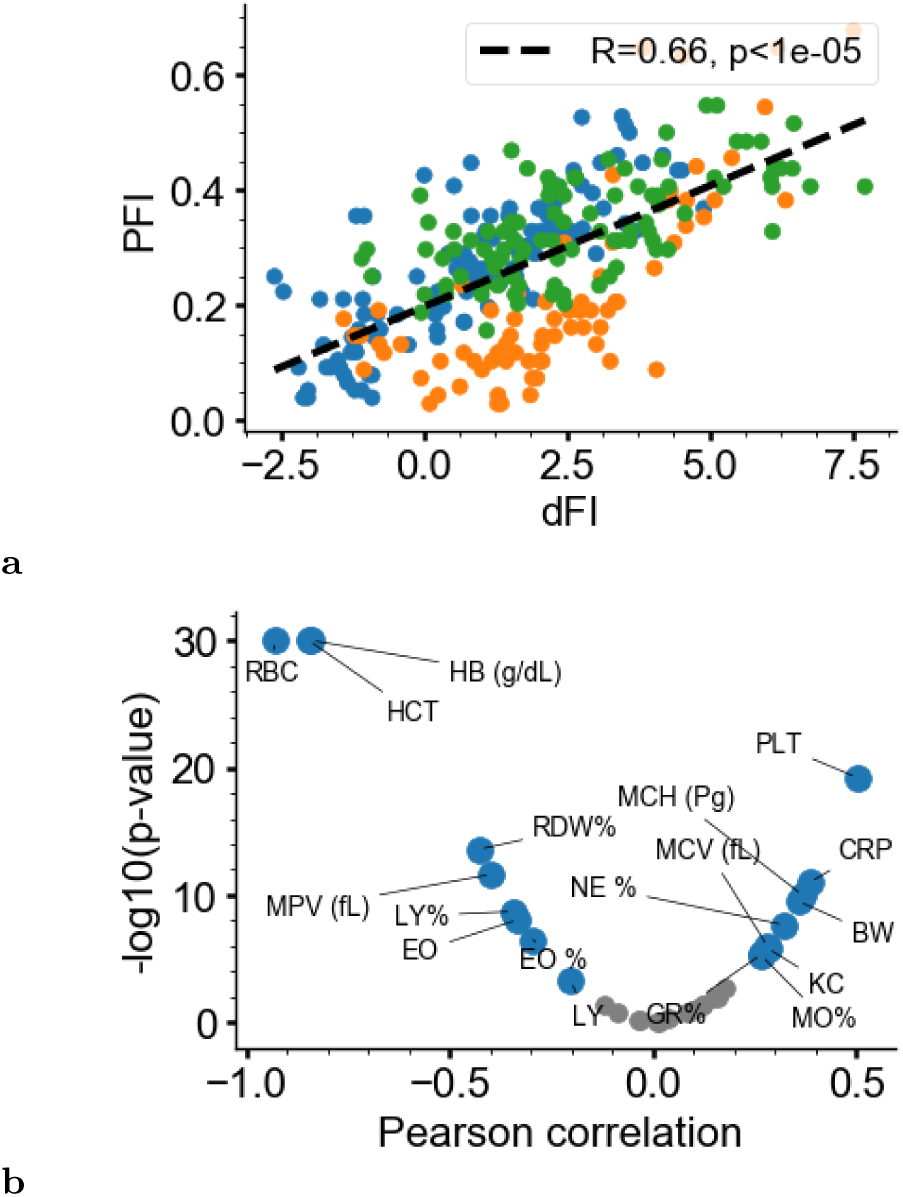
Correlation of dFI with the physiological frailty index (PFI) **a** and the extended set of phenotype measures **b** in the test datasets MA0071 and MA0072. Features with correlation above and below significance level 0.001 are shown with grey and blue circles, respectively. The most significant correlations (excluded dFI components) were between dFI and -reactive protein (CRP), red cell distribution width (RDW), body weight (BW) and murine chemokine CXCL1 (KC).

**FIG. 5:**
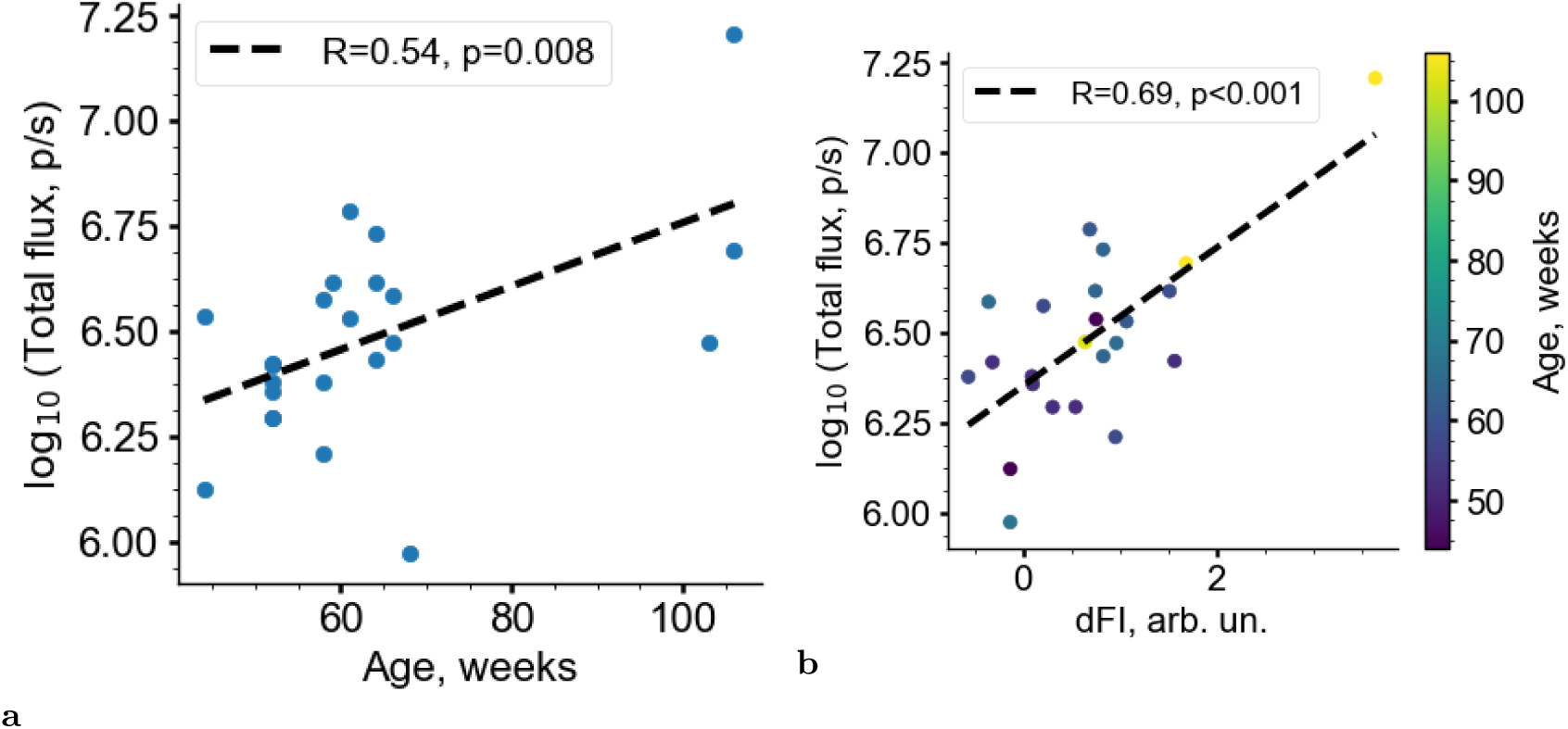
Total flux (TF) in log scale representing p16-dependent luciferase reporter activity as a quantitative indicator of senescent cells: statistically significant correlations with age (**a**) and with dFI (**b**) in old mice (> 50 weeks).

**FIG. 6:**
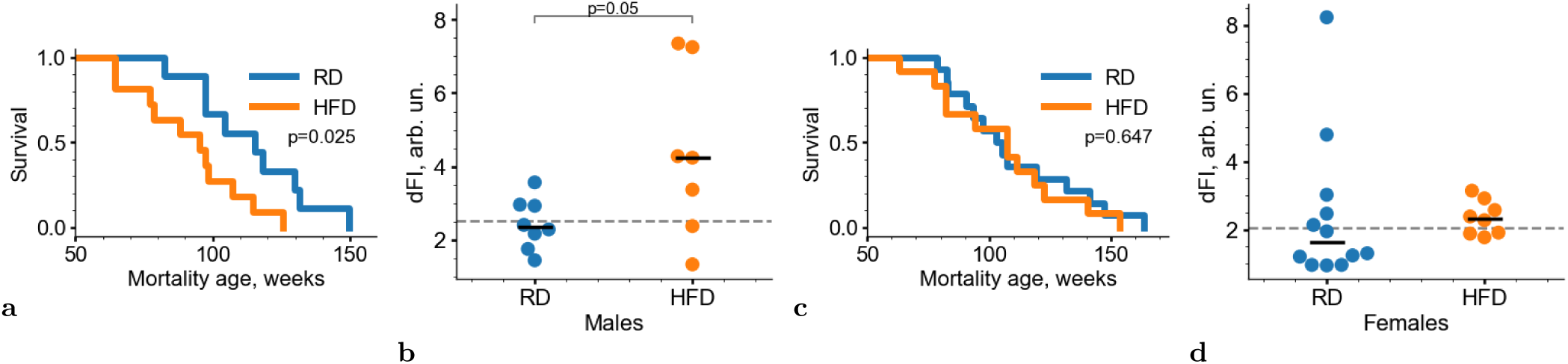
dFI responds to the lifespan-modifying effect of high-fat diet (HFD) feeding. (**a, c**) Kaplan-Meier survival curves showing that long-term (26 weeks) HFD feeding significantly reduces the lifespan of male (**a**) (p = 0.025, log rank test), but not female (**c**) (*p* = 0.6, log rank test) mice in comparison with regular diet (RD) feeding. (**b, d**) dFI values measured late in life (at week 78) for male (**b**) and female (**d**) mice fed with RD or HFD. Individual animals are represented by dots, with the horizontal bar indicating the group mean value. The horizontal dashed line shows the mean value for animals from both groups. dFI was significantly higher in males with HFD vs RD (*p* = 0.05, Student’s t-test), but there was no significant difference between HFD and RD groups of female mice.

**FIG. 7:**
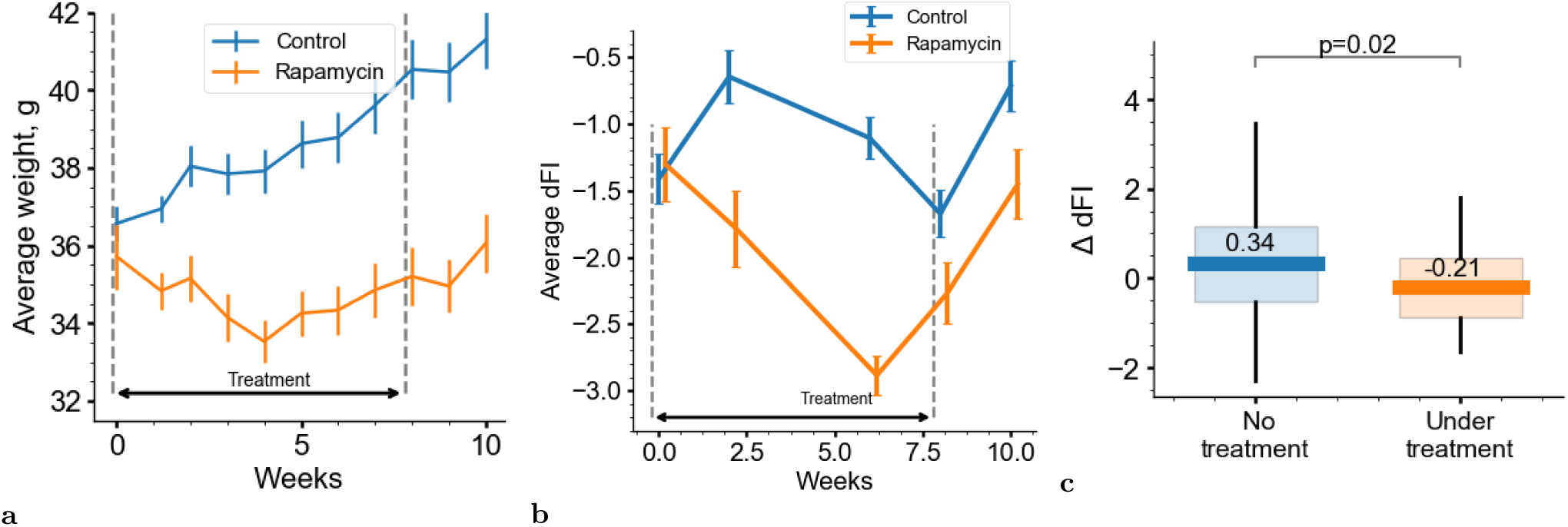
Effects of 8 week-long rapamycin treatment on body weight and dFI. Body weight (**a**) and dFI (**b**) were measured every one and two weeks, respectively. All of the data are presented as mean±SEM (*n* = 12 mice/group); (**c**) Change of dFI between two consecutive measurements when no treatment was given (blue box, includes both control and rapamycin group after withdrawal) or treatment was given (orange box, rapamycin group during treatment period). The change in dFI was signicantly lower under treatment (*p* = 0.02, Student’s two-tailed t-test).

**FIG. 8:**
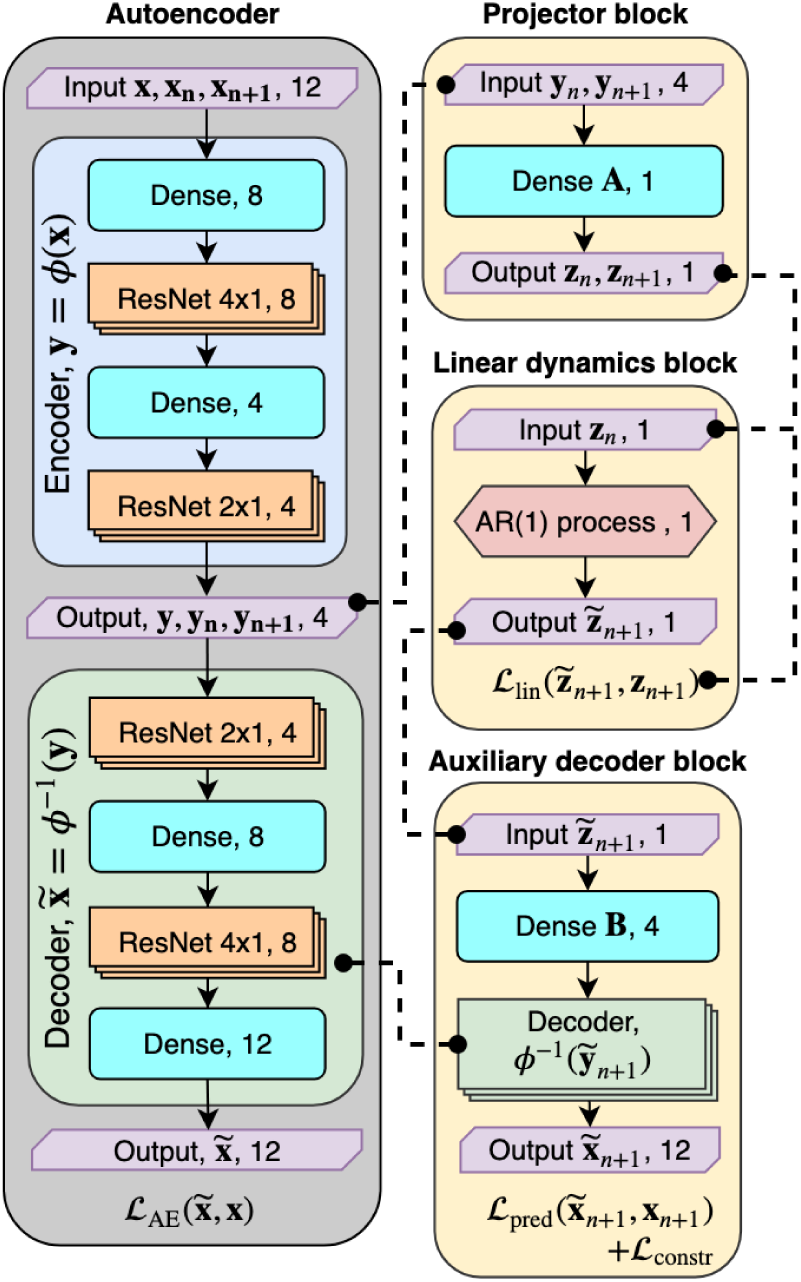
Network architecture of nonlinear auto-encoder (AE) with the embedded modes identication. The network is composed of AE, projector, linear dynamics and auxiliary decoder blocks. The AE block encodes input of 12 CBC parameters to the 4dimensional vector and then reconstructs back the original input. The AE consists of fully connected dense layers and residual network blocks (ResNet), which adds nonlinear rectication transformations. The AE is trained simultaneously on cross-sectional and longitudinal datasets. The projector block takes a 4−dimensional vector as an input and transforms it to a scalar *z*, which we refer to as dFI. During training, a pair of vectors is fed to the inputs: one *y*_*n*_ for the present state of the system and one *y*_*n*_ + 1 for the future state. The linear dynamics block solves the equation of rst-order autoregressive processes and predicts the future state *z*_*n*_ + 1. The auxiliary decoder block reconstructs the original 12−dimension CBC vector from the output of the linear dynamics block utilizing the decoder from the AE block.

The longitudinal slice of MPD has only a few hundred of specimen with serial measurements. The combined AE-AR approach adopted here let us maximize the number of mice used for training of the complete model. Thus, we were able to use all the available samples, including both the cross-sectional and the longitudinal segments of the MPD, in the AE arm of the algorithm to produce the highest quality low-dimensional representation of the data.

The performance of the models was validated in test datasets (see Material and Methods and Table S3), which were completely excluded from fitting. The test datasets were obtained from independent experiments by collecting CBC samples from cohorts of NIH Swiss mice of different age and sex (dataset MA0071), cohort NIH Swiss male mice observed for 15 months (dataset MA0072) and cohorts of naive male and female NIH Swiss mice that were humanely euthanized after reaching approved experimental endpoints (dataset MA0073).

We estimated the reconstruction error of the AE by calculation of the root mean squared error (RMSE) and coefficient of determination *R*^2^ for each CBC feature in training and test sets (Tables S4 and S5). The average RMSE in the test set was 228.8 with *R*^2^ = 0.54; in the training set, RMSE was 106.4 and *R*^2^ = 0.77. The best reconstruction was achieved for hematocrit (*R*^2^ = 0.94), red blood cells (*R*^2^ = 0.92) and lymphocytes (*R*^2^ = 0.87); the worst results were for mean corpuscular hemoglobin concentration (*R*^2^ = −0.9) and platelets (*R*^2^ = −0.12) in the test set.

Simultaneously with the AE, we trained the network to fit the longitudinal slice of MPD (including fully-grown animals at ages from 26 to 104 weeks with a sampling interval of Δ*t* = 26 weeks) to the solution of the linearized (*g* = 0) version of Eq.1,

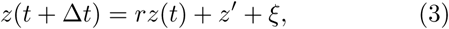

where *z* is the best possible linear combination of AE bottle-neck features. The state *z* is the output of the algorithm, the estimation of dFI (refer to the detailed description of the artificial neural network architecture behind the AE-AR algorithm in Fig. 8). The constants *r* = exp(*α*Δ*t*) ≈ 1, *z*′, and *ξ*, which are the best fit values of the autoregression coefficient, the constant shift, and the error of the fit (the combination of the system’s noise and measurement errors), respectively.

Performance of the AR model was demonstrated by plotting the autocorrelations between dFI values measured along aging trajectories of the same mice at age points separated by 14 and 28 weeks in the test dataset MA0072 (Fig. 2). Remarkably, the correlations (Pearson’s *r* = 0.71 (*p* < 0.001) and *r* = 0.70 (*p* < 0.001)) of the age-adjusted dFI persisted over the time lags of 14 and 28 weeks. The dFI auto-correlations were better than the autocorrelations of the first PC score *z*_0_ for the same mice, see Fig. S2; the corresponding Pearsons correlation values were *r* = 0.58 (*p* < 0.001) and *r* = 0.66 (*p* = 0.002) for 14- and 28-week time lags, correspondingly.

A semi-quantitative view of hierarchical clustering of CBC features co-variances in the test dataset produced groups of features associated with the immune system (white blood cell counts and the related quantities), metabolic rate/oxygen consumption (red blood cell counts and hemoglobin concentrations), and an apparently independent subsystem formed by platelets (see Fig. S4).

dFI was associated with animal age in both the training and test (see Fig. S3 and 3, respectively). As expected from the qualitative solution of Eq.1, dFI increased up to the age corresponding to the average animal lifespan (approximately 100 weeks in our case). We performed an exponential fit in the form of Eq.2 on the data from the test datasets (excluding animals that lived longer than the strains average lifespan and animals at the end of their life from the dataset MA0073). The calculation returned dFI growth exponent of *α* = 0.022 per week. This estimate is somewhat smaller than (but still of the same order as) the expected Gompertz mortality acceleration rate of 0.037 per week [24] for the SWR/J strain.

Saturation of the dFI beyond the average lifespan in the training and test datasets revealed a limiting value that is apparently incompatible with the animals’ survival. This possibility can be highlighted by plotting the dFI ranges from a separate cohort of “unhealthy” mice from MA0073 experiment, representing the animals scheduled for euthanasia under lab requirements (Fig. 3, red dots).

The long autocorrelation time of dFI together with its exponential growth at a rate compatible with the mortality acceleration rate are indicators of the association between dFI and mortality. This was further supported by the Spearman’s rank correlation between the dFI values and the order of the death events within mice in cohorts of same age and sex (Table II). We obtained significant correlations for dFI and remaining lifespan for all cohorts. Importantly, the age- and sex-adjusted dFI predicted remaining lifespan better than a naive PC score *z*_0_ from the linear analysis.

**TABLE II:**
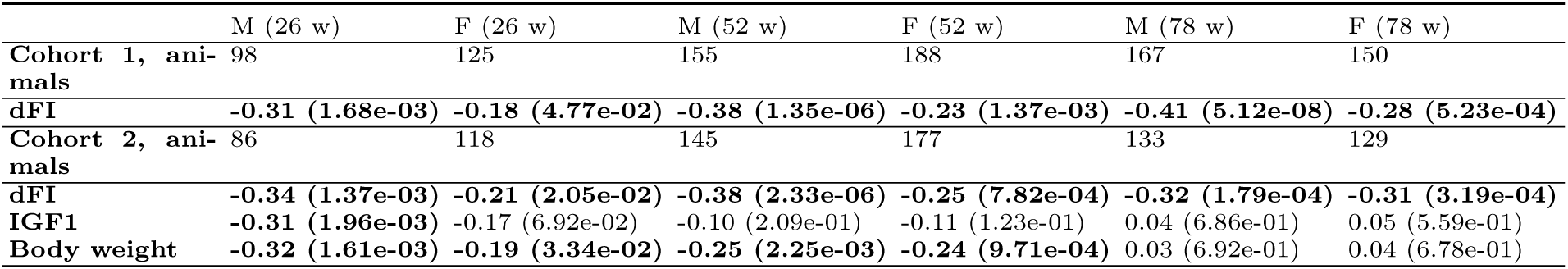
Spearman’s rank-order correlation values and the corresponding p-values (in parentheses) for dFI with lifespan. Analysis is shown for two cohorts: Cohort 1 includes all animals with mortality data, Cohort 2 includes the subset of animals from Cohort 1 for which IGF1 measurements were available. Significant correlations (*p* < 0.05) are highlighted in bold.

The dFI predicted remaining lifespan later in life better than body weight or insulin-like growth factor 1 (IGF1) serum level, which were previously shown to be associated with mortality in [25] and [26]. As pointed out in [26] and checked here, the concentration of IGF1 in serum was significantly associated with lifespan (*r* = −0.28, *p* = 0.008) only in one cohort of younger, 26-week old male mice. According to [25] and our calculations, mouse body weight is better associated with mortality, again, in the youngest animals at the ages of 26 and 52 weeks.

### D. dFI and hallmarks of aging

To further validate dFI as an age biomarker, we examined its association with physiological frailty index (PFI), a quantitative measure of aging and frailty established previously [19]. dFI and PFI were found to be strongly correlated (Pearson’s *r* = 0.64, *p* < 0.001), see Fig. 4**a**. PFI is a composite frailty score and depends on CBC measures for its determination. PFI is also influenced by changes in more traditional measures of frailty, such as grip strength, cardiovascular health, inflammation markers, etc. Remarkably, the correlation between PFI and dFI remained significant after adjustment for sex and age (Pearson’s *r* = 0.54, *p* < 0.001).

As illustrated in Figs. S4 and S6, we observed that the dFI was significantly associated with an extended set of CBC features across independent functional subsystems (most notably, but not limited to, myeloid cell lineage). The correlation between dFI and myeloid cell features was less profound in the training set, involving multiple strains (see Fig. S6). The correlation coefficient is a measure of response of dFI to individual CBC features variation and is different (sometimes even of opposite sign) in various mouse strains, see Fig. S7. The variation of the associations of individual features and dFI would be a significant challenge to a linear model and is a demonstration of the non-linear character of the autoencoder.

The dFI was strongly associated with red blood cell distribution width (RDW) and body weight (Fig. 4**b**), known predictors of frailty in both mice [15] and humans [27, 28]. dFI was also strongly associated with levels of C-reactive protein (CRP, *r* = 0.39, *p* < 0.001) and the murine chemokine CXCL1 (KC, *r* = 0.28, *p* < 0.001), both of which are known markers of systemic inflammation and mortality [29–31].

Aging is associated with an increasing burden of senescent cells [32, 33], widely considered to be a source of chronic sterile systemic inflammation, “inflammaging” [34]. Senescent cells are commonly detected in vivo as a population of p16/Ink4a-positive cells accumulated with age recognized by the activity of p16/Ink4a promoter-driven reporters [35]. We utilized earlier described hemizygous p16/Ink4a reporter mice with one p16/Ink4a allele knocked in with firefly luciferase cDNA [36]. Fig. 5**a** shows the correlation between animal age and presence of senescent cells, as measured by the flux from p16/Ink4a promoter-driven luciferase activity (*r* = 0.54, *p* = 0.01). The correlation of this SC proxy (total luciferase flux) with dFI was even stronger (*r* = 0.69, *p* < 0.001; Fig. 5**b**).

### E. dFI reflects lifespan-modulating interventions

Having established the association between dFI and remaining lifespan in the MPD, we next tested its predictive power by evaluating the response of dFI to life-long interventions known to affect the lifespan of mice. In the data from [19], male mice that were fed a high-fat diet (HFD) instead of a regular diet (RD) beginning at 50 weeks of age had significantly reduced lifespans (Fig. 6**a**) and also showed a significant increase in average dFI measured at week 78 (*p* = 0.05, Student’s two-tailed t-test; Fig. 6**b**) in comparison to control RD-fed males. In contrast, HFD feeding of female mice had no effect on either lifespan or average dFI (Figs. 6**c** and 6**d**). Thus, dFI appeared to be a good predictor of gender-dependent differences in organismal aging response to HFD, the underlying reasons for which remain to be explained.

We also tested the response of dFI to a short lifespan-extending condition: treatment with rapamycin [37, 38]. Here we present the results of an experiment with 60-week-old male mice treated with rapamycin daily at a dose of 12 mg/kg for 8 weeks or, in the control group, vehicle on the same schedule. The cohort of 24 60-week old C57BL/6 male mice was divided into treatment and control groups using a stratified randomization technique to produce indistinguishable distributions of dFI values.

Body weights were measured every week and increased as expected in the control group (Fig. 7**a**). In contrast, body weight in the rapamycin-treated group stayed approximately constant near the initial value throughout the observation period of 10 weeks. A lower body weight is typical for rapamycin-treated mice in comparison to control group [19, 39]. In order to generate dFI values for the mice in this experiment, blood samples were collected from each animal for CBC measurements every two weeks (Fig. 7**b**).

The longitudinal character of sampling in the experiment let us use the autoregression analysis to detect the effects of the drug on the dynamics of dFI in the course of the experiment. Whenever a non-random force (that is the effect of the drug) is present in Eq.1, the jump in dFI between any of consequent measurements from the same animal should satisfy modified Eq.3:

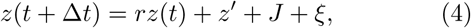

where *J* is the accumulated effect of the drug along the aging trajectory. The time intervals between the subsequent measurements are very small, *α*Δ*t* ≪ 1 and hence the autoregression coefficient *r* ≈ 1. We therefore expected to identify the effect of rapamycin by comparing the distributions of the dFI increments between the measurements.

We indeed observed the dFI jumps that were significantly different depending on whether rapamycin treatment was present between the dFI measurements both in the control and the treated groups, see Fig. 7**c** (*p* = 0.02, Student’s two-tailed test). These results support the possibility of using longitudinal dFI measurements to detect effects of life-extending therapeutics over much shorter times that what could be done based on the appearance of evident changes in frailty or longevity.

## III. DISCUSSION

We introduced a novel way of using deep artificial neuronal networks to train biomarkers of age and frailty from big biomedical data involving longitudinal measurements, i.e. multiple samples of the same animals collected along the aging trajectories. We exemplified the approach with the discovery and characterization of a novel biomarker of aging in mice, the dynamic frailty index (dFI), from conventional and automated measurements of Complete Blood Counts (CBC) and trained from the data from Mouse Phenome Database (MPD).

We started with linear dimensionality reduction using the principal component analysis (PCA), which has a long list of applications in biomarkers of aging research [40, 41]. As expected, we observed that the variance of CBC features in MPD is dominated by a cluster of features closely associated with the first PC score; none of the other PC scores correlated with age. Hence, the data suggests that aging in mice can be explained by the dynamics of a single (latent) variable that is a single organism-level quantity and a natural indicator of the progress of aging (i.e., a biomarker of aging).

The associations of slow organism state dynamics with the first principle component score is a hallmark of criticality, that is the situation whenever a system’s dynamics occurs in the vicinity of a tipping (or critical) point, separating the stable and the unstable regime [42]. Gene regulatory networks of most species are tuned to criticality [42]. In [18] we proposed that aging corresponds to the unstable regime, when small deviations of the organism state from its initial position get amplified exponentially. The first principal component score is then an approximation to the order parameter, herein referred to as dFI, that is corresponding to the unstable phase and having the meaning of the total number of the regulatory errors accumulated in the course of life of the animal [43].

The order-parameter is a generalization of a concept originally introduced in the Ginzburg-sLandau theory in order to describe phase-transitions in thermodynamics [44]. The order parameter concept was further generalized by Haken to the “enslaving-principle” saying that next to the critical point the dynamics of fast-relaxing (stable) components of a system is completely determined by the ‘slow’ dynamics of only a few ‘order-parameters’ (often variables associated with unstable modes) [17]. The dFI identified in connection with the dynamics of the order parameter is then not a mere machine learning tool for specific predictions, but a fundamental macroscopic property of the aging organism as a non-equilibrium system.

PCA belongs to the class of unsupervised learning algorithms, such that the model does not require any labels such as chronological age or the remaining lifespan for its training. It is therefore remarkable, although expected from a large corpus of previous works, that the first principle components are associated with age and the remaining lifespan of the animals. However, the abilities of linear rank reduction techniques, such as PCA, to recover accurate dynamic description of aging is limited for the following reasons. First, there are no reasons to believe that the effects of non-linear interactions between different dynamic subsystems are small. That is why the result of such a procedure can not be expected to perform well in different biological contexts (strains, laboratory conditions, or therapeutic interventions such as drugs).

Second, biological measurements are often noisy, and hence, simple techniques lacking efficient regularization may fail to reconstruct the latent variables space correctly unless a prohibitively large number of samples is obtained [45]. Finally, the association of the first principal component with the order parameter and hence the biomarker of aging in the form of dFI is only an approximate statement. Fundamentally, there is no way to identify the dynamics of the system from the data, that does not include the dynamics itself in the form of multiple measurements of the same organism along the aging trajectory.

To compensate for the drawbacks of PCA, we employed an artificial neuron network, a combination of a deep denoising auto-encoder (AE) and an auto-regressive (AR) model. The AE part of the algorithm is a non-linear generalization of PCA and was used to compress the correlated and necessarily noisy biological measurements into a compact set of latent variables, a low-dimensional representation of the organism state.

The AR-arm of the network is nothing else but the best possible prediction of a future state of the same animal from the current measurements in such a way that the collective variable inferred by the model is a directly interpretable and physiologically relevant feature, the dynamic frailty index (dFI). The approach is a computational metaphor for the analytical model behind identification of the order parameters associated with the organism-level regulatory network instability from [18]. The neural network applied here was inspired by deep rank-reduction architectures, recently used for characterization and interpretation of numerical solutions of large non-linear dynamical systems [46, 47].

dFI increases exponentially with age and is associated with remaining lifespan. It is therefore a natural quantitative measure of aging drift and hence may be used as a biomarker of age. Remarkably, it appears that blood parameter data alone can define biological age with a degree of accuracy comparable to that of the best previously described biomarkers of aging e.g., DNA methylation-based clock [5–8] or physiological frailty index [19]. This may reflect a key role for aging of hematopoietic tissue in determining aging of the whole organism, a concept that is intuitively acceptable given the universal systemic physiological function of blood.

As an alternative explanation, age-dependent changes in blood parameters may be secondary events induced by aging of the remainder of the organism (i.e., various solid tissues). However, accumulated experimental evidence argues against this. In fact, there are multiple reports demonstrating rejuvenating effects of young hematopoietic system on old animals delivered either by bone marrow transplantation or by parabiosis (reviewed in ref. [48]). Moreover, restoration of mouse hematopoiesis through transplantation of HSCs from young vs old donors clearly demonstrated that aged HSCs cannot be rejuvenated by the environment of a young body [49]. Also, the interpretation of age dependence of HSC-derived features as secondary effects of aging would face formal difficulties, since the dynamics of such factors should exhibit shorter, in fact at least twice shorter, doubling times than the dFI and the mortality rate doubling times.

A peculiar result of our analysis is that our data strongly point towards myeloid lineage that provides much more accurate predictors of biological age than lymphoid lineage parameters. This is counterintuitive since aging is generally accepted to be associated with the well-documented general decline in immunity known as an immunosenescence [50–52], the phenomenon illustrated by the reduced efficiency of vaccination [53] and increased frequency and lethality of infectious diseases and cancer in older organisms [54]. Nevertheless, there is strong experimental evidence that supports and provides a mechanistic explanation for our finding that myeloid parameters weigh more heavily than lymphoid ones as biological age indicators. In a comprehensive study of the epigenetic mechanisms of HSC aging, Beerman et al. [49]. described age-dependent epigenetic reprogramming that leads to a significant shift towards myeloid lineage differentiation of the progeny of aged HSCs [49, 55, 56]. This shift is driven by specific changes in methylation of the DNA of HSCs that occur during mouse aging. Surprisingly, these changes in methylation, which alter gene expression, do not occur in the part of the genome that controls HSC phenotype, but rather modify DNA regions encoding genes that control downstream differentiation stages. Remarkably, the pattern of DNA methylation changes associated with aging of HSCs seems to represent the same process that was previously described as a DNA methylation-based clock [5, 49], and therefore, may be part of the same epigenetically controlled fundamental aging mechanism. Another factor that could diminish the impact of lymphoid lineage-related parameters as biological age markers is the reactive nature of this branch of hematopoiesis, which serves to rapidly respond to sporadic events such as viral or bacterial infection, wounding, and other types of stress requiring an emergency response usually in the form of acute inflammation. Since the time of occurrence of such events is unpredictable, age-associated changes may be masked by the noise coming from large-scale age-unrelated fluctuations in the lymphoid compartment.

These observations do not mean that the blood is the single determinant of aging (otherwise, biological age would be 100% defined by the age of HSCs), but at least place it among the major drivers of the process and provide an explanation for our success in reliably determining biological age from blood test data. Rather, the identification of aging with the dynamics of a single organism state variable, dFI, suggests a cross-talk in the form of continuous interactions between the organism components. dFI, hence emerges as a feature characterizing the organism as a whole, rather than representing a property of any particular subsystem.

The cooperative character of aging in the model implies that there is no specific subsystems tracking time or age in an animal. The age-dependent chances appear in a self-consistent manner by strong non-linear interactions between physiological compartments. Formally, this is expressed by representing the aging organism as an autonomous (or time-invariant) dynamical system having no designated subsystem for tracking time. Accordingly, we expected no physiological indices may depend on age of the animals explicitly, only implicitly via dependence on the collective variable, dFI. That is why, we believe, the analysis of dFI properties revealed that in addition to the trivial dependence on CBC features, which were directly involved in dFI calculation, the dFI was strongly correlated with certain measures of frailty, also known as hallmarks of aging. These include grip strength, body weight, RDW, and markers of inflammation such as CRP and KC (IL-8). dFI also correlated well with p16-luciferase flux, a proxy for the number of senescent cells in aged mice. We observed a very high degree of concordance between the dFI and the physiological frailty index (PFI), which is a combination of a much wider range of analyses than CBC, including physical fitness, cardiovascular health and biochemistry.

The dFI increased at a characteristic doubling rate of 0.022 per week, that is, in line with our theoretical prediction, comparable with the mortality rate doubling time in the species. Also, in the cross-sectional dataset the dFI saturated at a limiting value at the age corresponding to the average lifespan in the group. However, we observed that the dFI ceiling corresponds to the dFI levels in cohorts of animals scheduled for euthanasia due to morbid conditions under current laboratory protocols, which is as close to death as animals could possibly be in a modern laboratory. Therefore, we conclude that further dFI increments are incompatible with survival. It is thus dynamics of the organism state defining the unconstrained growth of dFI fueled by the dynamic instability of the organism state is the ultimate cause of death in aging mice. In [18], we explained that the exponential dFI acceleration is a signature of the linear dynamics in the weak coupling limit. At the maximum dFI level, the inevitably present non-linear effects take over and further evolution of the organism state occurs on much shorter time scales and lead to a complete disintegration of the organism.

The effects of non-linearity can be neglected nearly always in the course of the life of an animal, if the dimensionless parameter expressing the animal lifespan in units of the mortality rate doubling time is small. Given the observed dFI doubling rates, we infer that the corresponding ratio is of the order of two, which is hardly large, and hence, non-linear corrections to the dynamics of the order parameter, dFI, should not be very small. Therefore, our linear AR model is only a reasonable approximation. We therefore believe that better performing dFI variants could be obtained by allowing for higher rank AR models, possibly including the effects of mode coupling with dFI.

The deep artificial neural network applied here also belongs to the class of unsupervised algorithms. It is remarkable that we used neither the remaining lifespan nor even the chronological age of the animals to infer dFI. This was possible, in principle, since by having a very specific model of the aging process, we were able to use longitudinal aging trajectories of individual animals for training. Due to the ability to obtain meaningful description of aging in the data without health or lifespan labels, the proposed method should be particularly useful for analysis of large longitudinal datasets from recently introduced sensors (such as wearable devices) often without any clinical and/or survival follow-up information available.

Aging manifests itself as slow deviations of the organism state from its initial state and can be tracked by measuring dFI. Our analysis shows that that the underlying organism state regulatory network in mice is dynamically unstable, and hence the organism state cannot relax to any equilibrium value after a perturbation. Formally this is expressed by strong auto-correlations of dFI over extended periods of time. It is therefore likely that the effects of short treatments should persist until the end of life, whereas the effects of such treatments could be detected in short experiments involving longitudinal dFI measurements over a few months’ time.

The dynamic character of dFI implies that most of the organism state changes associated with aging are in fact reversible. We therefore expect that further investigation of the longitudinal dynamics of physiological state variables and the associated biomarkers of aging and frailty could eventually lead to cost- and time-efficient clinical trials of upcoming anti-aging therapeutics.

## IV. MATERIALS AND METHODS

### A. Datasets

The training data set of CBC features was prepared from the nine data sources available in the Mouse Phenome Database (MPD) [13, 14]. List of the included sources is presented in Table S2 together with a statistic on animal number group by sex and age cohorts. Our model was trained using the best overlap of available CBC features from all sources. The final list contained 12 CBC features: granulocytes differential (GR%), granulocytes count (GR), hemoglobin (HB), hematocrit (HCT%), lymphocyte differential (LY%), lymphocyte count (LY), mean corpuscular hemoglobin content (MCH), mean hemoglobin concentration (MCHC), mean corpuscular volume (MCV), platelet count (PLT), red blood cell count (RBC) and white blood cell count (WBC). In the case of data source had no granulocytes measurements, it was retrieved using formulas:

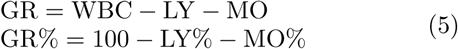

All animals with the missing data were excluded from the training.

The list of all abbreviations is shown in Table S1.

### B. Animals

Four-to-five week-old NIH Swiss male and female mice were obtained from Charles River Laboratories (Wilmington, MA) and were allowed to age within the Roswell Park Comprehensive Cancer Center (RPCCC) animal facility. Blood samples were obtained at different ages as part of creating of the Physiological Frailty Index (PFI) as previously described (REF). p16/INK4a-LUC mice (p16-Luc) were obtained from the N. Sharpless laboratory at the University of North Carolina (Chapel Hill, NC). All animals were housed under 12:12 light:dark conditions (12 hours of light followed by 12 hours of darkness) at the Laboratory Animal Shared Resource at RPCCC. All animal experiments were approved by the Institutional Animal Care and Use Committee of Roswell Park Cancer Institute.

Dataset MA0071 was built in a cross-sectional experiment using male and female NIH Swiss mice. Blood was collected from male mice by cardiac puncture at 26 (*n* = 20), 64 (*n* = 20), 78 (*n* = 20), 92 (*n* = 20), and 136 (*n* = 8) week old mice. Female age groups were represented by 30 (*n* = 20), 56 (*n* = 20), 68 (*n* = 20), 82 (*n* = 20), 95 (*n* = 20), 108 (*n* = 20), and 136 (*n* = 8) weeks of age.

Dataset MA0072 was obtained from a longitudinal experiment. Blood samples were collected through saphenous vein from male NIH Swiss mice at 66 (*n* = 30), 81 (*n* = 24), 94 (*n* = 22), 109 (*n* = 18), and 130 (*n* = 11) weeks of age.

Dataset MA0073 includes blood samples collected from 97 male and 127 female mice of different ages when animals reached approved experimental endpoints and require humane euthanasia. Whole blood cell analysis was performed in 20 ul of blood using Hemavet 950 Analyzer (Drew Scientific) according to manufacturer’s protocol. For rapamycin treatment experiment 60-weeks-old C57BL/6J male mice were obtained from Jackson Laboratories (USA). Rapamycin was purchased from LC Laboratories (MA, USA). Rapamycin was administered daily at 12 mg/kg via oral gavage for 8 weeks. Control group was treated with vehicle (5% Tween-80, 5% PEG-400, 3% DMSO).

### C. In vivo bioluminescence imaging

Bioluminescence imaging was performed using an IVIS Spectrum imaging system (Caliper LifeSciences, Inc, Waltham, MA). p16/Ink4a-Luc+/- female mice were injected. intraperitoneally with D-Luciferin (150 mg/kg, Gold Biotechnology), 3 minutes later anesthetized with isoflurane and imaged using a 20-second integration time and medium binning. Data were quantified as the sum of photon flux recorded from both sides of each mouse using Living Image software (Perkin Elmer, Waltham, MA.).

### D. Dimensionality reduction with PCA

Principal component analysis (PCA) was performed with Python [57] and Scikit-learn package [58]. First, we applied PCA transformation on the entire training dataset. However, the principal components were dominated by the difference of mice strains. Animals of the same strains were clustered on the plot of the first principal component against the second one. We removed strain difference by subtracting mean values of CBC features calculated for the earliest age available for the selected strain from values of CBC features of all animals for this strain. For the simplicity we restricted our analysis to 30 strains, which were presented in the Peters4 dataset.

### E. Statistical analysis of mortality data

The death records for animals linked with the MPD dataset Peters4 were also available in MPD as the dataset named Yuan2 [59]. The Spearman’s rank correlation test was performed with Python and SciPy package [60]. The analysis was performed for two cohorts of mice. The first cohort included all animals from the Peters4 dataset with mortality data from Yuan2. The second cohort included animals from the Peters4 dataset with the measurements of body weight and IGF1 serum level taken from MPD dataset named Yuan1 [26].

### F. Neural network structure

The neural network was designed to handle a specific problem: the disbalance of samples with longitudinal and cross-sectional measurements. As inputs, the network has three 12-dimensional vectors: one for the cross-sectional dataset, and two others for the longitudinal dataset corresponding to the present state and future state of a sample. Inputs pass through the encoder part of the auto-encoder block and then split up (see Fig. 8). Cross-sectional samples are directed to the decoder part, while longitudinal samples in the compressed representation are passed for the training autoregression part. Such data flow allows the auto-encoder to be deeper and train without overfitting by using more samples from a larger cross-sectional dataset. The auto-encoder has the architecture of a linear stack of fully connected dense layers and residual network blocks (ResNet) [61]. Dense layers have a trainable weight matrix **W**, bias vector **b**, and linear activation function by default. The ResNet block, shown in Fig. 9, is a stack of two dense layers with an activation function of rectified linear unit (ReLU)[62], input and output are linked by applying element-wise addition. ResNet blocks add nonlinear rectification transformations to the original input, helping to learn non-linear transformations. To prevent overfitting, we applied *L*2 regularization of factor 0.01 to model weights **W**.

**FIG. 9:**
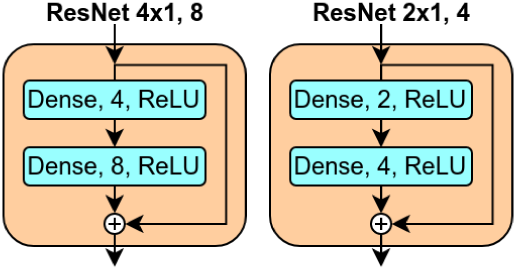
Schematic of residual network (ResNet) blocks. These consist of two fully connected dense layers with activation function of rectied linear unit (ReLU). Input and output are interconnected by applying element-wise addition.

## V. ACKNOWLEDGEMENTS

We thank Norman Sharpless for generously providing p16-Luc reporter mice. This work was supported in part by National Cancer Institute (NCI) grant P30 CA016056 for use of Roswell Park Comprehensive Cancer Centers Laboratory Animal and Translational Imaging Shared Resources (MPA and AVG) and by a contract from Genome Protection, Inc. (MPA and AVG).

