## Supplementary Tables and Figures for "Identification of a blood test-based biomarker of aging through deep learning of aging trajectories in large phenotypic datasets of mice"

### Supplementary information

Konstantin Avchaciov et al.

#### S1 Supplementary tables

| Abbreviation | Full name | Units |
|---|---|---|
| BW | body weight | g |
| CRP | plasma concentration of C-reactive protein | pg/mL |
| Dias | diastolic blood pressure | mmHg |
| EO and EO% | eosinophil number and differential | K/ul and % |
| Flow | tail blood flow rate | cm/s |
| GLU | plasma glucose concentration | pg/mL |
| GR and GR% | granulocytes number and differential | K/ul and % |
| HB | hemoglobin | g/dL |
| HCT | hematocrit | % |
| HR | heart beat rate | bpm |
| IGF1 | plasma concentration of IGF1 | ng/mL |
| INS | plasma insulin concentration | pg/mL |
| KC | plasma concentration of pro-inflammatory cytokine KC | pg/mL |
| LY and LY% | lymphocyte number and differential | K/ul and % |
| MCH | mean corpuscular hemoglobin | Pg |
| MCHC | mean corpuscular hemoglobin concentration | g/dL |
| MCV | mean corpuscular volume | fL |
| MO and MO% | monocyte number and differential | K/ul and % |
| MPV | mean platelet volume | fL |
| NE and NE% | neutrophil number and differential | K/ul and % |
| PLT | platelet count | K/ul |
| RBC | red blood cell counts | M/uL |
| RDW | red cell distribution width | % |
| Sys | systolic blood pressure | mmHg |
| TG | plasma triglycerides concentration | pg/mL |
| Volume | tail blood volume | cm/s |
| WBC | white blood cell count | K/ul |

Table S1: List of abbreviations.

| MPD dataset | Sex | # of strains | Age, weeks | # of animals | Training autoen-coder | Training autoregres-sion |
|---|---|---|---|---|---|---|
| CGDpheno1 | F | 71 | 11 | 711 | + | |
| | M | 72 | 11 | 670 | + | |
| CGDpheno3 | F | 62 | 7 | 321 | + | |
| | M | 61 | 7 | 321 | + | |
| Jaxpheno4 | F | 11 | 8 | 205 | + | |
| | | | 16 | 155 | + | |
| | M | 11 | 8 | 203 | + | |
| | | | 16 | 149 | + | |
| Justice2 | F | 16 | 14 | 202 | + | |
| | M | 16 | 14 | 196 | + | |
| Lake1 | F | 23 | 7 | 186 | + | |
| | M | 23 | 7 | 188 | + | |
| Peters1 | F | 43 | 11 | 762 | + | |
| | M | 43 | 11 | 713 | + | |
| Peters2 | F | 28 | 11 | 222 | + | |
| | M | 28 | 11 | 227 | + | |
| Peters4 | F | 30 | 26 | 234 | + | + |
| | | | 52 | 232 | + | + |
| | | | 78 | 195 | + | + |
| | | | 104 | 74 | + | |
| | M | 30 | 26 | 221 | + | + |
| | | | 52 | 223 | + | + |
| | | | 78 | 229 | + | + |
| | | | 104 | 82 | + | |
| Svenson3 | F | 28 | 7 | 348 | + | |
| | M | 28 | 7 | 347 | + | |

Table S2: Description of MPD datasets used for training.

| Study | Sex | Strain | Age, weeks | # of animals |
|---|---|---|---|---|
| MA0071 | F | NIH Swiss | 30 | 20 |
| | | | 56 | 20 |
| | | | 68 | 20 |
| | | | 82 | 19 |
| | | | 95 | 19 |
| | | | 108 | 20 |
| | | | 136 | 8 |
| | M | NIH Swiss | 26 | 20 |
| | | | 64 | 18 |
| | | | 78 | 19 |
| | | | 92 | 15 |
| | | | 132 | 6 |
| MA0072 | M | NIH Swiss | 66 | 29 |
| | | | 81 | 23 |
| | | | 94 | 21 |
| | | | 109 | 16 |
| | | | 130 | 9 |
| MA0073 | M | NIH Swiss | 66-160 | 97 |
| | F | NIH Swiss | 66-160 | 127 |

Table S3: Description of test datasets.

| | gr % | gr (k/ul) | hb (g/dl) | hct % | ly % | ly (k/ul) |
|---|---|---|---|---|---|---|
| CGDpheno1 | 3.54 (0.89) | 0.81 (-0.12) | 0.70 (0.74) | 0.59 (0.98) | 3.68 (0.89) | 0.94 (0.89) |
| CGDpheno3 | 1.89 (0.82) | 0.22 (0.62) | 0.37 (0.70) | 0.32 (0.98) | 2.00 (0.81) | 0.43 (0.94) |
| Jaxpheno4 | 3.66 (0.94) | 0.25 (0.66) | 0.66 (0.55) | 0.58 (0.96) | 3.71 (0.94) | 0.28 (0.94) |
| Justice2 | 2.29 (0.79) | 0.24 (0.64) | 0.42 (0.70) | 0.36 (0.98) | 2.11 (0.85) | 0.39 (0.96) |
| Lake1 | 2.71 (0.21) | 0.46 (-0.93) | 0.55 (0.55) | 0.46 (0.97) | 3.42 (-0.12) | 0.74 (0.85) |
| Peters1 | 3.05 (0.86) | 0.52 (0.61) | 0.57 (0.74) | 0.46 (0.99) | 3.05 (0.86) | 0.76 (0.91) |
| Peters2 | 2.40 (0.67) | 0.24 (0.61) | 0.36 (0.80) | 0.40 (0.97) | 2.50 (0.68) | 0.60 (0.93) |
| Peters4 | 3.95 (0.93) | 0.87 (0.74) | 0.67 (0.80) | 0.66 (0.98) | 3.99 (0.93) | 0.81 (0.94) |
| Svenson3 | 2.57 (0.84) | 0.33 (0.55) | 0.62 (0.54) | 0.64 (0.99) | 2.79 (0.83) | 0.71 (0.92) |
| Total | 3.23 (0.92) | 0.60 (0.69) | 0.60 (0.80) | 0.54 (0.99) | 3.33 (0.93) | 0.72 (0.94) |

Table continuation:

| | mch (pg) | mchc (g/dl) | mcv(fl) | plt (k/ul) | rbc (m/ul) | wbc (k/ul) |
|---|---|---|---|---|---|---|
| CGDpheno1 | 0.48 (0.80) | 1.67 (0.37) | 1.17 (0.83) | 355 (-0.07) | 0.23 (0.94) | 1.88 (0.65) |
| CGDpheno3 | 0.27 (0.85) | 0.87 (0.48) | 0.56 (0.94) | 209 (-0.14) | 0.13 (0.94) | 0.73 (0.85) |
| Jaxpheno4 | 0.50 (0.72) | 1.54 (0.18) | 0.83 (0.90) | 234 (-0.12) | 0.17 (0.95) | 0.55 (0.83) |
| Justice2 | 0.32 (0.85) | 0.97 (-0.37) | 0.57 (0.92) | 220 (-0.11) | 0.15 (0.95) | 0.65 (0.91) |
| Lake1 | 0.47 (0.68) | 1.64 (0.25) | 0.94 (0.39) | 146 (0.12) | 0.17 (0.89) | 1.47 (0.50) |
| Peters1 | 0.43 (0.81) | 1.52 (0.34) | 0.95 (0.89) | 376 (0.04) | 0.20 (0.95) | 1.31 (0.81) |
| Peters2 | 0.28 (0.76) | 0.98 (0.27) | 0.63 (0.89) | 203 (-0.29) | 0.14 (0.93) | 1.05 (0.83) |
| Peters4 | 0.43 (0.81) | 1.67 (0.34) | 1.15 (0.86) | 575 (0.02) | 0.30 (0.92) | 1.84 (0.78) |
| Svenson3 | 0.43 (0.71) | 1.41 (0.68) | 0.99 (0.91) | 197 (-0.13) | 0.17 (0.94) | 1.21 (0.80) |
| Total | 0.42 (0.81) | 1.48 (0.47) | 0.97 (0.89) | 369 (0.07) | 0.21 (0.95) | 1.43 (0.83) |

Table S4: Reconstruction error (root-mean-square error, RMSE) and coefficient of determination, $R^2$, of the autoencoder calculated for each CBC feature in the training set

| | gr % | gr (k/ul) | hb (g/dl) | hct % | ly % | ly (k/ul) |
|---|---|---|---|---|---|---|
| MA0071 | 7.56 (0.55) | 0.66 (0.50) | 0.96 (0.78) | 0.74 (0.99) | 6.06 (0.80) | 0.58 (0.69) |
| MA0072 | 5.72 (0.62) | 0.52 (0.72) | 0.76 (0.89) | 1.42 (0.96) | 4.64 (0.73) | 0.41 (0.95) |
| MA0073D | 9.94 (0.66) | 2.00 (0.38) | 1.44 (0.78) | 2.99 (0.91) | 7.58 (0.79) | 0.85 (0.88) |
| Total | 8.34 (0.68) | 1.37 (0.51) | 1.15 (0.82) | 2.07 (0.94) | 6.51 (0.84) | 0.68 (0.87) |

Table continuation:

| | mch (pg) | mchc (g/dl) | mcv(fl) | plt (k/ul) | rbc (m/ul) | wbc (k/ul) |
|---|---|---|---|---|---|---|
| MA0071 | 0.69 (-0.19) | 2.59 (-5.22) | 1.76 (0.38) | 498 (0.08) | 0.33 (0.94) | 2.12 (0.43) |
| MA0072 | 0.59 (0.67) | 2.09 (-0.34) | 1.20 (0.87) | 1127 (-1.03) | 0.26 (0.96) | 0.98 (0.87) |
| MA0073D | 0.91 (0.66) | 4.15 (-1.11) | 3.65 (0.56) | 823 (-0.09) | 0.68 (0.87) | 3.10 (0.63) |
| Total | 0.77 (0.67) | 3.26 (-0.90) | 2.65 (0.59) | 792 (-0.12) | 0.50 (0.92) | 2.43 (0.64) |

Table S5: Reconstruction error (root-mean-square error, RMSE) and coefficient of determination, $R^2$, of the autoencoder calculated for each CBC feature in the test set

#### S2 Supplementary figures

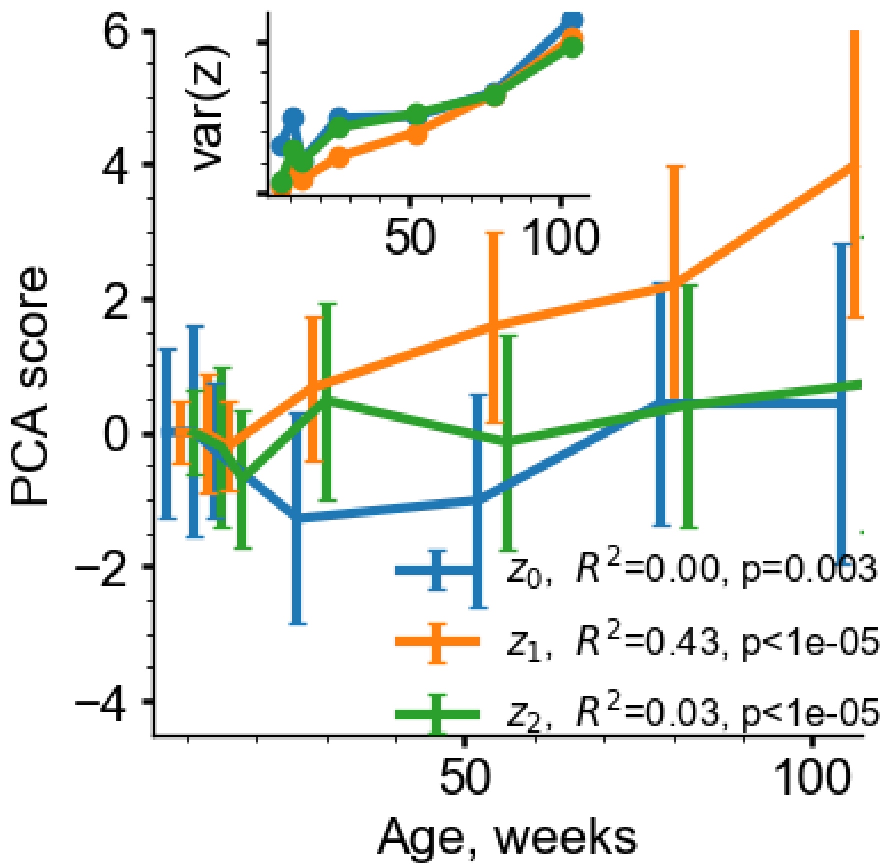


Figure S1: Principal Component Analysis (PCA) of the MPD data (including young animals). The graphs represent the average of the PC scores in subsequent age groups. The inset shows that the variance for all PC scores increase with age.

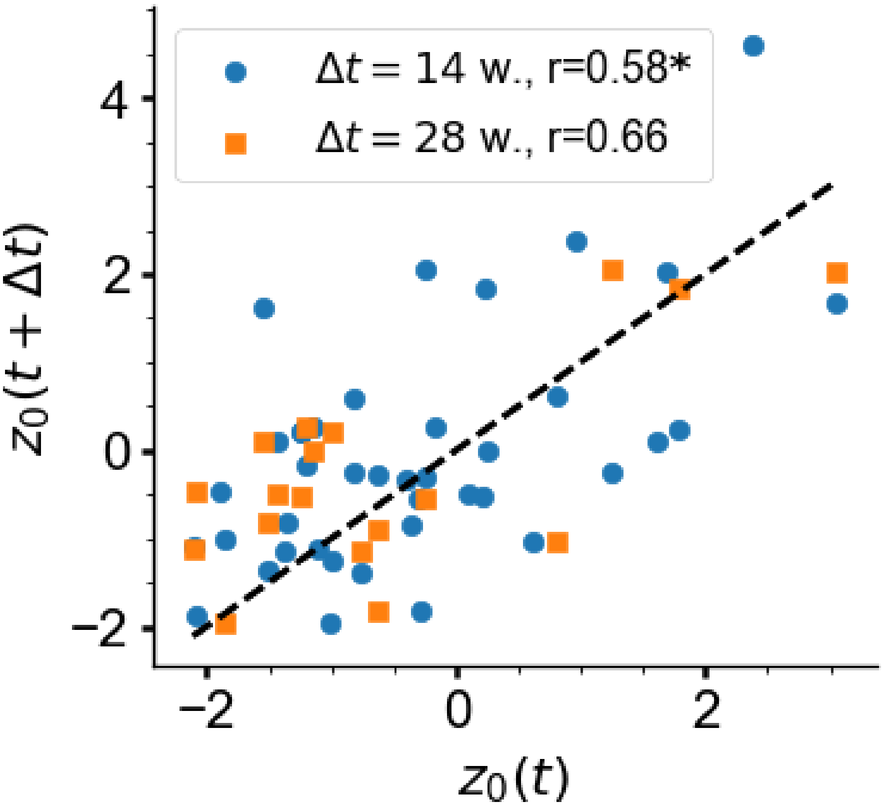


Figure S2: Auto-correlation properties of age-adjusted rst PC score ($z_0$) in the test longitudinal dataset with sampling intervals $\Delta t$ of 14 (blue circles) and 28 (orange squares) weeks. $*$ marks statistically significant correlations, $p < 0.001$.

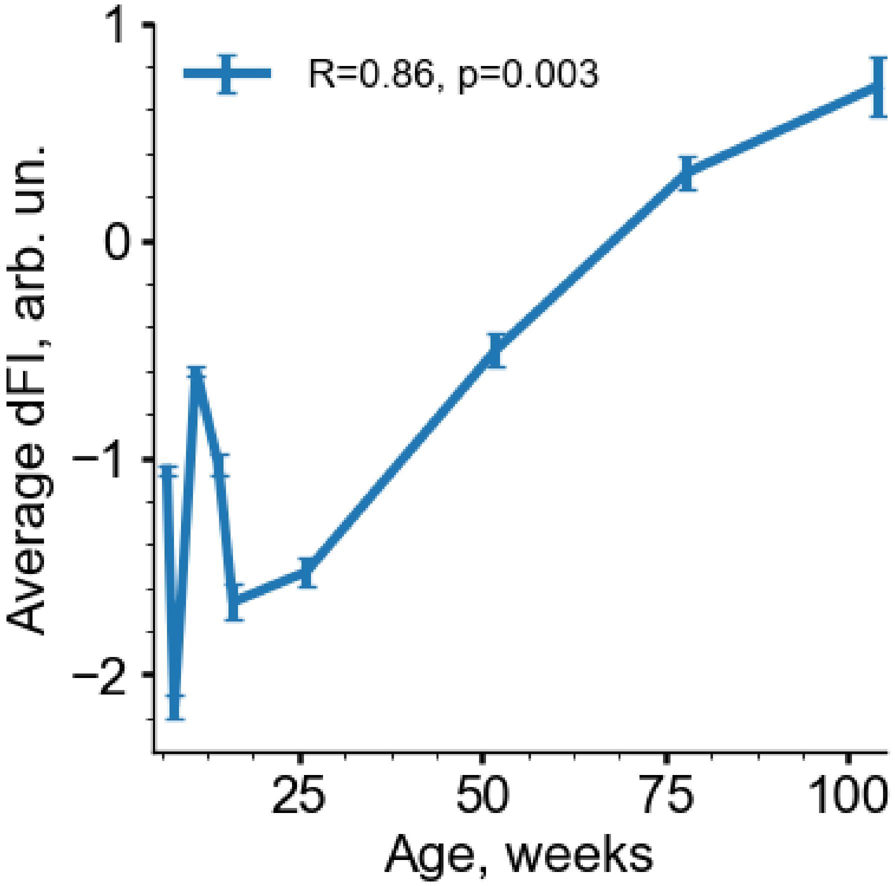


Figure S3: The growth of age cohort average dFI with age in the training dataset.

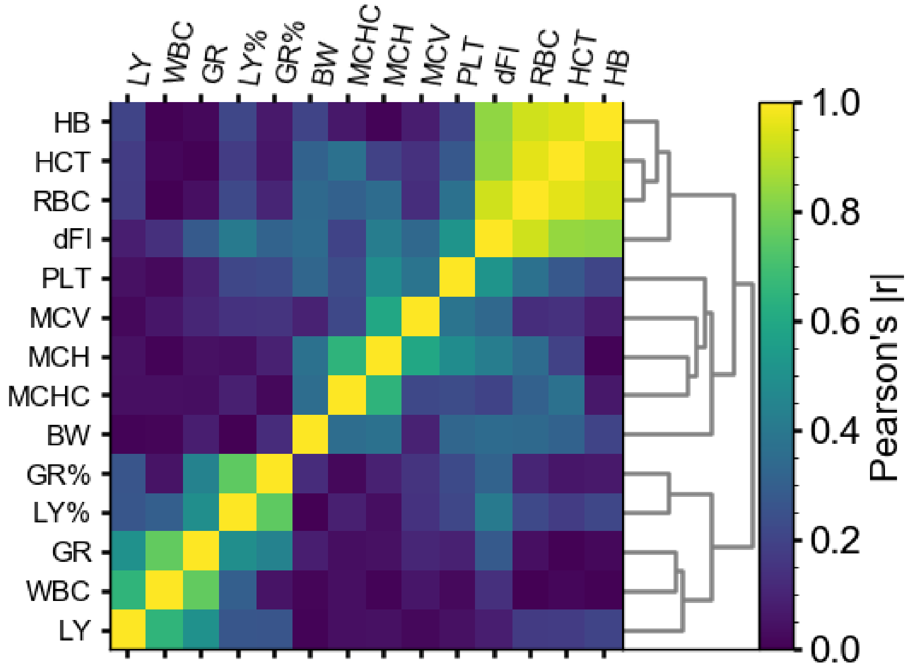


Figure S4: Clustering of CBC features and dFI score in the test dataset. The colors represent the Pearson's correlation coefficient (absolute value) as indicated by the scale on the right side of the figure.

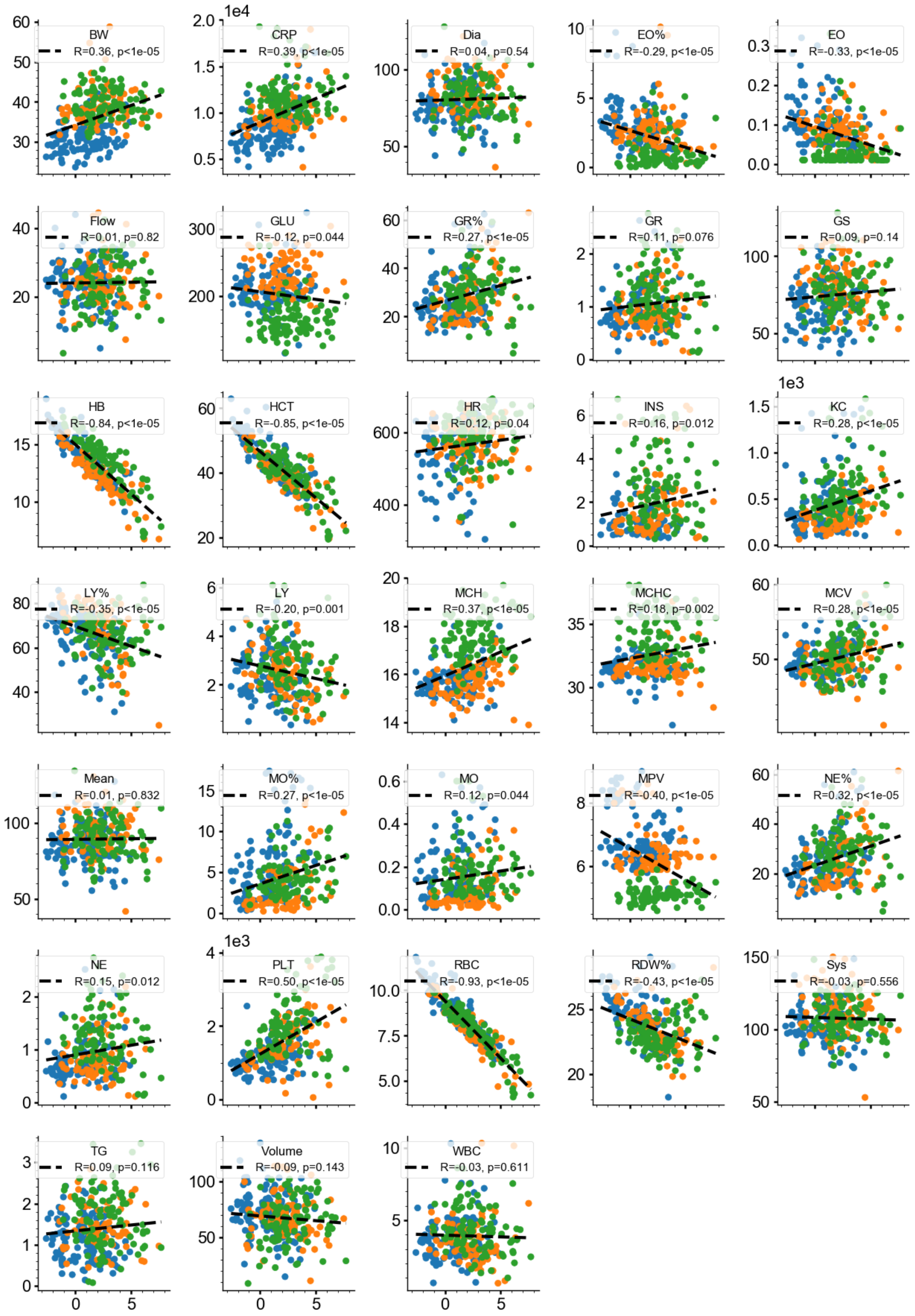


Figure S5: Correlations between dFI and other biological markers. Colors represent datasets females in MA0071 (blue), males in MA0071 (orange) and males in MA0072 (green).

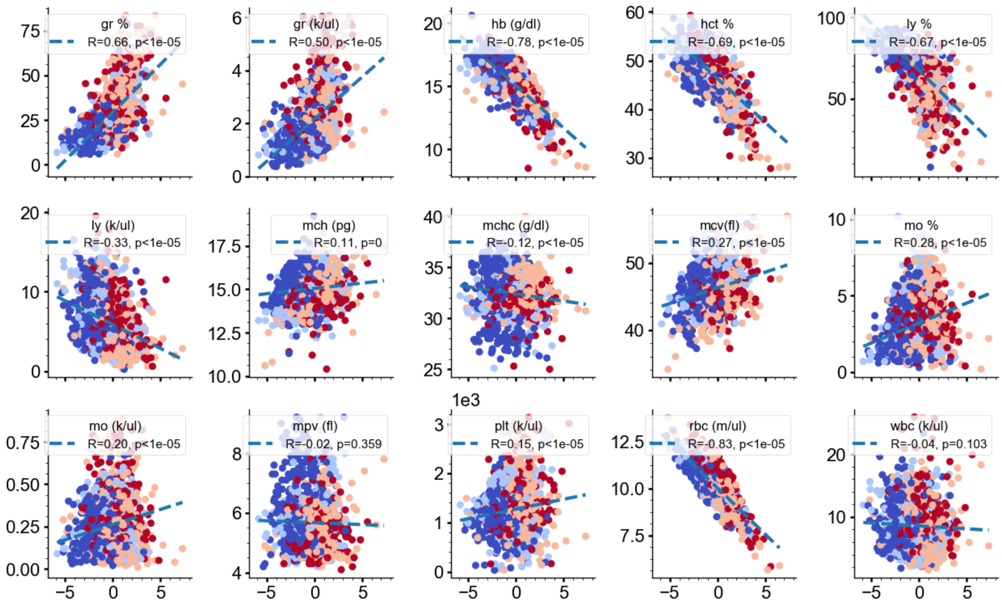


Figure S6: Correlations between dFI and CBC parameters in the *Peters4* dataset. Colors from blue to red represent age of animals, where blue is age of 26 weeks and red is age of 104 weeks.

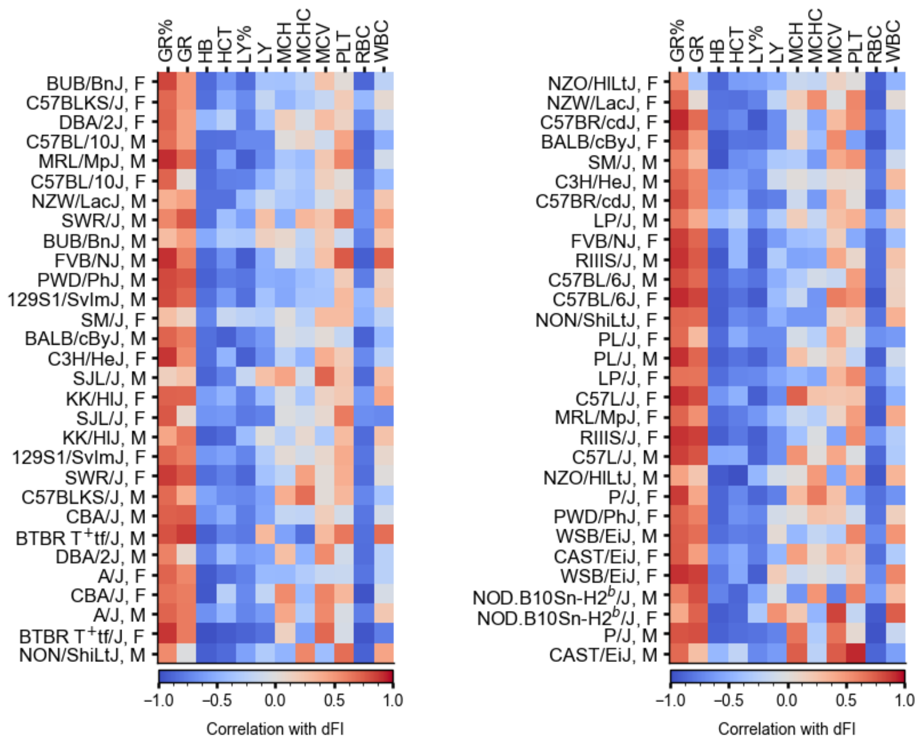


Figure S7: Correlations between dFI and CBC parameters in the *Peters4* dataset shown for cohort of mice of same strain and sex.
